# Long vs. short read sequencing for microbial ecology of sedimentary environments: a case study from Lake Arnon, Switzerland

**DOI:** 10.1101/2025.07.20.665787

**Authors:** Thomas Camille, Aliisa K. Laakkonen, Deborah R. Rast, Kremer Katrina, Max Shore, Vogel Hendrik

## Abstract

The subsurface biosphere remains poorly characterized, with many resident microorganisms uncultured and lacking genomic references. Despite the growing accessibility of shotgun metagenomics, 16S rRNA gene sequencing remains a standard tool for microbial community profiling, often relying on sequence similarity to reference databases such as SILVA to infer taxonomy and potential function. However, in environments with low biomass and high proportions of unknown lineages, such as deeper sedimentary environments, the accuracy of these inferences and our ability to capture rare taxa remain uncertain. A better inference of these rare taxa may now be possible with the advent of accurate long-read applications that have recently become available.

Here we provide a comparison of long-read (PacBio) and short-read (Illumina NextSeq) 16S rRNA approaches for microbial communities from a sediment core of Lake Arnon (Switzerland). We compared community composition in environmental samples and mock controls to evaluate the strengths and limitations of each method. While sequencing technology significantly influenced observed community structure, sediment depth had an even stronger effect. Taxonomic profiles were broadly consistent across methods for most bacterial groups, but archaeal diversity was underrepresented in the long-read data, likely due to primer mismatch. When detected, long-read sequencing offered more accurate taxonomic resolution, often down to the species level, enabling better inference of metabolic potential. Beta diversity patterns were similar at broad taxonomic levels between methods, though more detailed metrics such as species contributions to beta diversity (SCBD) and co-occurrence networks showed enhanced resolution and specificity in long-read datasets. Our results highlight the critical importance of primer design, in particular for capturing archaeal taxa that play important roles in the deep biosphere. With improved primer coverage and continued cost reductions, long-read sequencing holds strong potential for advancing our understanding of subsurface microbial identity, structure and function.

## 1. Introduction

The subsurface biosphere remains one of the least understood habitats on Earth, with a significant portion of its microbial diversity yet to be explored (1–4). Many of the organisms inhabiting this environment have never been cultured, and their genomes remain uncharacterized, making it challenging to infer their ecological roles, although progress has been made on this end recently (see reviews by Escudero et al., 2018 or Orsi, 2018 for examples).

In this context, high-throughput sequencing of the 16S rRNA gene is the most common method applied in microbial community profiling, allowing researchers to classify taxa based on sequence similarity to known databases (7–9). Advancements in sequencing technologies and bioinformatics have significantly improved the accuracy of taxonomic assignments, while metagenomics shotgun sequencing approaches have provided a more comprehensive view of this "dark biosphere" (e.g. Baker et al., 2016; Rinke et al., 2013). However, metabarcoding remains a widely used and cost-effective method for characterizing biodiversity in environments where high-quality DNA is scarce. Despite its accessibility and efficiency, it is still subject to inherent biases and limitations that will impact the resolution and completeness of microbial community assessment (12,13). Among the factors influencing the obtained microbial diversity in subsurface environments are DNA extraction methods (14–16), primer choices (17–19), sequencing approaches including read length and sequencing depth (20–22), and post-sequencing data treatment and analyses (including chimera treatment, ASV/OTU approach and database choice) (23–25).

The characteristics of subsurface biosphere studies include low cell number, poor DNA quality, low number of culturable representatives, and issues of interfering substances during DNA processing (e.g. acidity of the medium, humic substances and inhibitors affecting amplification, binding of DNA to clay) (23,26). Standard Short Reads Illumina 16S rRNA gene-based approaches can lead to incomplete capture of the full extent of ecological potentials but are, based on assessments, still considered appropriate to estimate biodiversity in diverse systems, even when comparing with shotgun approaches (27,28). When dealing with rare taxa, low-biomass samples, or highly divergent microbial communities, the challenge is compounded by sequencing biases, primer mismatches, and database limitations, all of which can affect the resolution of taxonomic assignments, particularly for archaeal communities, which are often underrepresented in reference datasets (Seitz et al., 2019; Orsi et al., 2020). However, some studies actually showed the potential for full-length 16S rRNA metabarcoding in revealing specific functions and interactions, even in sediments or methane-rich environments where *Archaea* were important actors (29,30). Exploring this potential in subsurface environments is therefore of interest. To further address these challenges, we conducted a comparative study of microbial communities in the sedimentary environment of Lake Arnon, an alpine lake in Switzerland. We employed two distinct metabarcoding approaches for 16S rRNA: PacBio HiFi long-read sequencing, which provides full-length 16S rRNA gene sequences with improved taxonomic resolution, and the most common Illumina NextSeq short-read sequencing, focusing on the V3-V4 region, which offers high-throughput, with a deep sequencing coverage. We set the other parameters (DNA extraction method, sequence post-processing, taxonomy database) to avoid interference of other parameters.

By applying both methods to community controls and natural sediment samples from the first ca 50 cm of sediments, we aim to evaluate their respective strengths and limitations in microbial ecology research. Specifically, we compared their ability to:

1. Accurately classify microbial taxa, particularly those from poorly characterized lineages.
2. Capture sediment-specific microbial diversity, including low-abundance archaeal taxa.
3. Assess how different sequencing approaches influence community composition and taxonomic affiliations.

By comparing long-read and short-read sequencing approaches in the context of Lake Arnon sediments, we contribute to the ongoing discussion about the best strategies for capturing microbial diversity in challenging environments.

## 2. Lake setting

Lake Arnon is a reservoir lake located in the Bernese Oberland region of Switzerland, at an elevation of approximately 1,540 meters above sea level. Originally filling a basin carved during the last glaciations, the natural lake was expanded and transformed into a reservoir in 1942.

The sedimentation of Lake Arnon is characterized by (1) allochthonous sources: mineral substrates eroded from the surrounding slopes and organic material (residues from the surrounding vegetation made of alpine meadows and coniferous forests) from the catchment are accumulating in the basin, and by (2) autochthonous sources from in-situ primary production. The allochthonous input is mainly controlled by snowmelt and precipitation. In addition, the lake is surrounded by steep slopes, favoring landslides into the lake. (Fig. 1). The combination of these geological and hydrological factors shapes the sedimentary environment of Lake Arnon, potentially influencing the microbial communities inhabiting its lakebed. Additionally, the dam construction and maintenance, along with leisure activities on the lake may influence its overall biodiversity.

**Fig. 1.**
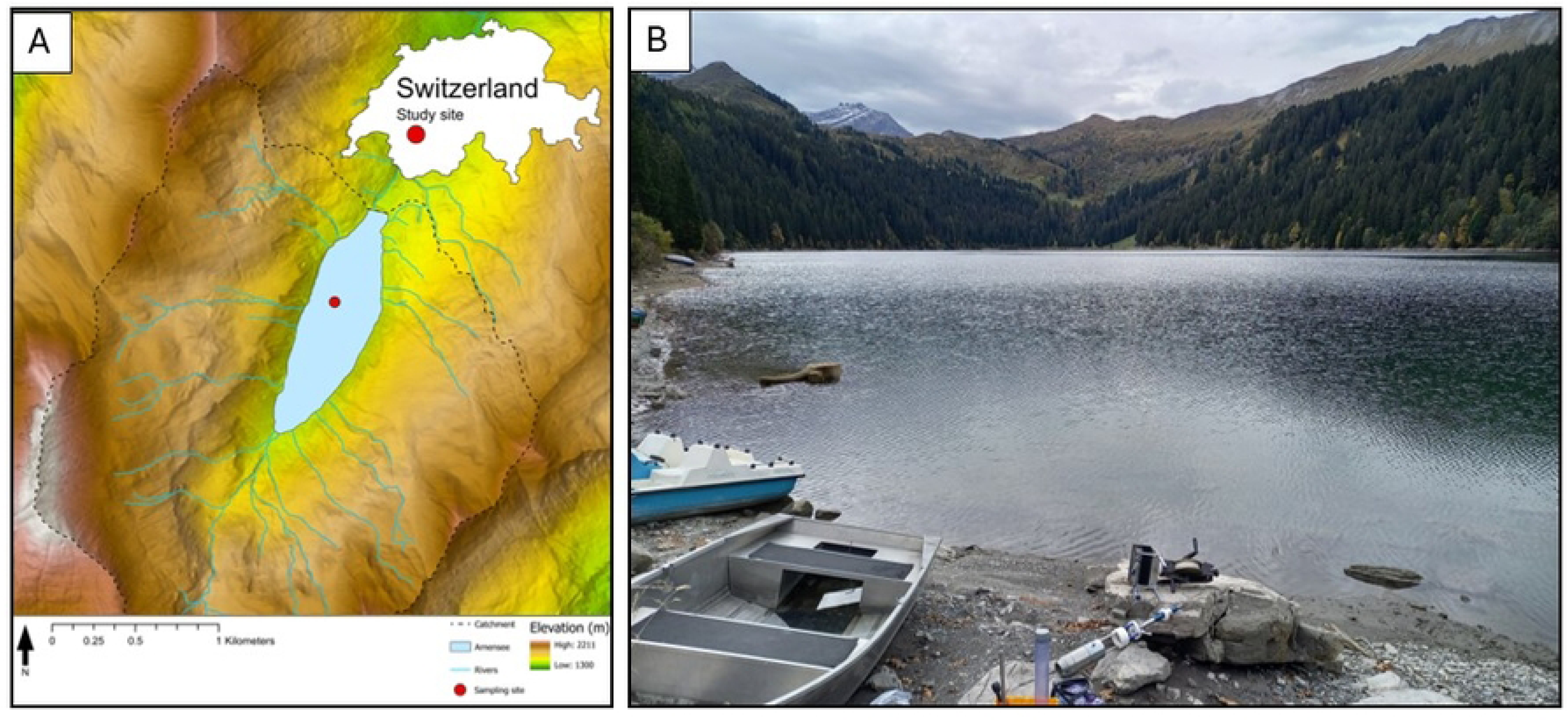
Description of study site. (A) Location of study site on Lake Arnon within the Swiss Alps. (B) Photograph of the lake and surrounding vegetated landscape, with coring material in fore front.

## 3. Methods

### 3.1. Lake sediment coring and sampling

A 55 cm core was obtained from the center of Lake Arnon (BE, Switzerland), at ca. 45 m water depth, using a gravity corer and a pre-pierced liner to allow sterile sampling for DNA. Pre-cut and autoclaved syringes were inserted in the pre-pierced holes of the liners (initially covered with tape) to sample at given depths of 1-2-3-4-5-6-7-8-9-10-15-20-25-30-40-50 cm. Wet sediment (1 cm^3^) was transferred from the syringes to a sterile cryotube filled with 3mL of Qiagen PowerProtect DNA/RNA preservative (Qiagen, Germany). The mud in contact with the liner/tape was discarded. The samples were then stored in a cool box for 3 hours (rest of the field day) and then transferred to -20°C freezer until extraction occurred.

### 3.2. DNA extraction

Samples were thawed, vortexed and 1 mL of mixed sediment and PowerProtect DNA/RNA preservative (approximately 300 to 500 mg of wet sediment) were transferred to bead tubes of the DNeasy PowerSoil Pro Kit (Qiagen, Germany). DNA was then extracted using the manufacturer guidelines, with vortexing using a Vortex-Mixer Genie 2 during 15 min at maximum speed and a final elution step in 50 uL.

### 3.3. Sequencing : long read and short reads

Prior to amplification of full-length bacterial 16S gene region V1-9 with barcoded primers, the starting DNA was assessed for quantity, quality and purity using a Qubit 4.0 flurometer (Qubit dsDNA HS Assay kit; Q32851, Thermo Fisher Scientific), an Advanced Analytical FEMTO Pulse instrument (Genomic DNA 165 kb Kit; FP-1002-0275, Agilent) and a Denovix DS-11 UV-Vis spectrophotometer, respectively. Thereafter, the PCR amplification of bacterial full-length 16S rRNA genes (V1—V9 regions) was done according to the PN 101-599-700 procedure and checklist document from PacBio

For the amplification of bacterial full-length 16S rRNA genes the following primers were used: 16S rRNA degenerate forward primer sequence* 5’ GCATC/barcode/AGRGTTYGATYMTGGCTCAG 3’, known as the 27F primer and 16S rRNA degenerate reverse primer sequence* 5’ GCATC/barcode/RGYTACCTTGTTACGACTT 3’ known as the 1492R primer barcode. Four controls (ctr) were always included, 2 negative controls (EB and MM) and Zymo_Com_ctrl (https://zymoresearch.eu/collections/zymobiomics-microbial-community-standards/products/zymobiomics-microbial-community-dna-standard), Zymo_Log_ctrl (https://zymoresearch.eu/products/zymobiomics-microbial-community-dna-standard-ii-log-distribution).

The barcoded amplicons were evaluated by using a Thermo Fisher Scientific Qubit 4.0 fluorometer with the Qubit dsDNA HS Assay Kit (Thermo Fisher Scientific, Q32854) and an Agilent Fragment Analyzer (Agilent) with a HS NGS Fragment Kit (Agilent, DNF-474), respectively. They were pooled equi-volume and a SMRT bell library was generated according to the Preparing multiplexed amplicon libraries using SMRTbell prep kit 3.0 procedure and checklist document PN 102-359-000 from PacBio following the Primer-indexed sample workflow. Library pool concentration and size were again assessed (Qubit and FEMTO Pulse as described above), respectively. Instructions in SMRT Link Sample Setup were followed to prepare the SMRTbell library for sequencing (PacBio SMRT Link v11.1). Shortly, PacBio Sequencing primers v3.1 and Sequel DNA Polymerase 2.1 were annealed and bound, respectively, to the DNA template libraries using a Sequel II Binding Kit 3.1 (PacBio Part number 102-333-400) and the complex was cleaned using SMRTbell clean-up beads. The libraries were loaded at an on-plate concentration of 110pM using adaptive loading, along with the use of Sequel II DNA internal control complex 3.1. SMRT sequencing was performed in CCS mode on the Sequel IIe with Sequel Sequencing kit 3.0, SMRT Cells 8M, a 2h pre-extension followed by a 10 h movie time and via PacBio SMRT Link v11.1. Thereafter, the CCS generation is performed on the Sequel IIe to generate highly accurate single molecule reads (HiFi reads) and the barcode demultiplexing workflow was run in SMRT Link v11.1.

For short-read sequencing, NGS libraries targeting the V3-V4 region of the 16S rRNA gene were constructed using the Quick-16S Plus NGS Library Prep Kit (V3-V4) (Zymo Research, D6420-PS1) according to the instruction manual version 1.2.1. Five ng of input DNA was used for each sample, a BIO-RAD CFX96 Real-time System and C1000 touch thermal cycler were used for the qPCR-based steps. Alongside the samples, two negative controls were included; one with the buffer used to dilute the samples and one with the PCR mastermix lacking any DNA template. Furthermore, a Microbial Community DNA Standard and a Microbial Community DNA Standard II (Log Distribution) were also processed (Zymo Research, ZymoBIOMICS D6306 & D6311, respectively). The final library pool was evaluated using Qubit and Agilent Fragment Analyzer; respectively. The library pool, including all controls, was sequenced 300 bp paired-end using a NextSeq1000/2000 P1 Reagents (600 cycles) kit (illumina, 20075294) on an illumina NextSeq 1000 instrument. The quality of the sequencing run was assessed using illumina Sequencing Analysis Viewer (illumina version 2.5.12) and all base call files were demultiplexed and converted into FASTQ files using illumina bcl2fastq conversion software v2.20.

All steps post DNA extraction to sequencing data generation and data utility were performed at the Next Generation Sequencing Platform, University of Bern, Switzerland

### 3.4. Data processing

PacBio HiFi reads were processed using the pb-16S-nf pipeline developed for full-length 16S rRNA amplicon data (https://github.com/PacificBiosciences/HiFi-16S-workflow). The analysis was run using both default and relaxed filtering settings to evaluate their impact on archaeal detection and diversity recovery. Key parameter adjustments included the VSEARCH identity threshold (default 0.97, tested values down to 0.7), maximum expected error (maxEE) (default 2, tested up to 5), min_asv_totalfreq (default 5, tested value 0), and min_asv_sample (default 1, tested value 0). Taxonomic assignment was performed using a Naïve Bayes classifier on the SILVA nr99 v138.1 database.

For both LR and SR approaches, samples were filtered and trimmed using cutadapt (31), assembled and checked for chimeras using dada2 (32) and further analyzed with phyloseq (33) and vegan (34) packages in R (R Core Team, 2013). Sample labelling was done as follow: SR or LR for Short Reads and Long Reads respectively, followed by the sample depth in the sediment, in cm. Samples SR-2, SR-3, SR-10 and LR-10 were discarded as they did not pass filtering steps (too low read number).

Microbial community composition was compared between datasets obtained using short-read (SR) and long-read (LR) sequencing approaches. To assess patterns of community variation, the data were collapsed at the phylum level based on taxonomic assignments. Rarefaction was applied to standardize sequencing depth by down-sampling the SR dataset to the minimum read count observed in the LR dataset. LR/SR ratio was calculated based on this rarefaction.

Non-metric multidimensional scaling (NMDS) based on Bray-Curtis dissimilarity was performed on relative abundance data using the metaMDS function in the vegan package. The influence of sequencing method and depth on community structure was evaluated using permutational multivariate analysis of variance (PERMANOVA) with 999 permutations (adonis2, vegan).

To further explore systematic differences in taxonomic composition between SR and LR datasets, log₂ fold changes in mean relative abundances were calculated at the phylum level. Phyla were grouped based on fold-change magnitude to highlight both strongly biased and consistently detected taxa across sequencing approaches. The archaeal classes were also reported in a barplot for SR. Relative abundance was mapped along depth for a number of taxa showing significant importance in our betadiversity analysis.

Beta diversity was assessed following the variance partitioning framework of Legendre and De Cáceres (2013), using a custom implementation of the beta.div() function. The analysis was based on a Hellinger-transformed ASV abundance matrix derived from the LR and SR dataset. Species contributions to beta diversity (SCBD) for each taxon were calculated and ranked, and the top 100 contributing ASVs were visualized using bar plots. Top 50 and 20 ASV were pooled by taxonomic affiliation in circular plots for better reading, for LR and SR respectively. Co-occurrence networks were constructed separately for LR and SR datasets using the igraph (v2.0.3) and Hmisc (v5.1-2) packages in R. To reduce noise and improve network interpretability, amplicon sequence variants (ASVs) were filtered to retain only those present in at least two samples. Pairwise Spearman’s rank correlations were computed, and edges were retained based on correlation coefficients > |0.7| and p-values < 0.01. For comparability, the number of ASVs in the SR dataset was rarefied to match that of LR (∼1200 ASVs). Community modules within each network were identified using a fast greedy modularity optimization algorithm. Node-level metrics and modular assignments were extracted using igraph and tibble. Networks were visualized in Gephi (v0.10.1) using the Force Atlas 2 layout, with node size scaled by degree and colors indicating either modular affiliation or taxonomic group (phylum level).

R scripts, metadata and sequences are available on the OSF page of the project https://osf.io/35hjr/ with DOI 10.17605/OSF.IO/35HJR. Sequences were deposited under SUB15450934 on NCBI, and a final accession number will be provided after revisions.

## 4. Results

### 4.1. Sequencing Diversity and Depth

The observed sequencing read counts range from approximately 608 to 966 reads for PacBio LR sequencing, compared to a broader span of 21,969 to 41,443 reads for Illumina SR (Fig. 2A-B). Notably, when sample LR-7 is excluded, the minimum read count shifts to sample 40 across both sequencing platforms, suggesting potential issues related to DNA quality and extraction efficiency in this sample. The highest read counts were observed in samples LR-9 for LR and SR-5 for SR. These differences in sequencing depth are mirrored in the Shannon diversity indices, which average 5.9 for LR and 10.1 for SR, with the lowest diversity values also observed in sample 40 (LR: 5.3, SR: 9.5). Other samples demonstrate more consistent Shannon values, falling within 6.1 to 6.5 for LR and 10.0 to 10.3 for SR (Fig. 2C). To further explore diversity variations and assess the impact of sequencing approach (LR vs. SR), we rarefied the Illumina short reads to the minimum PacBio read count of 600, analyzing how the Shannon index ratio (LR/SR) changes with sequencing depth. Post-rarefaction, the average Shannon index for LR reads was 5.61, with a variance of 0.06 and a standard deviation of 0.24. For SR reads, the average Shannon index stabilized at 6.37, with negligible variance and a standard deviation of 0.02. We used thus this rarefied approach in the Shannon index ratio (LR/SR), which exhibited notable variation, from 0.78 at 40 cm depth to 0.92 at 6 cm depth, and demonstrated a slight decreasing trend with increasing depth (Fig. 2D).

**Fig. 2.**
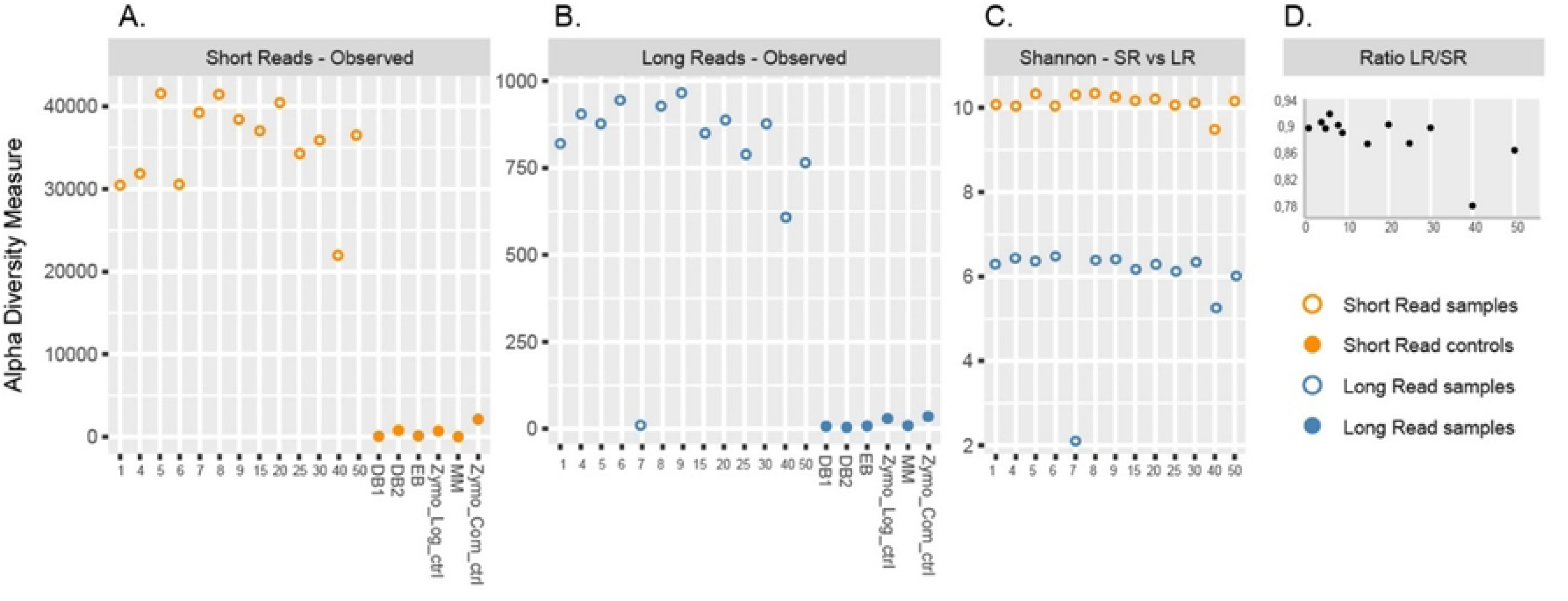
Alpha diversity metrics for 16S rRNA gene sequencing for lake Arnon samples and controls. (A) Richness for SR. (B) Richness for LR. (C) Shannon diversity index for SR and LR samples. (D) LR/SR diversity index per sample across depth

### 4.2. Taxonomic Affiliation Variations by Sequencing Method and Depth

Taxonomic affiliation analysis against the SILVA database revealed a similar composition throughout the samples, with notable differences between LR and SR sequencing approaches. Virtually all taxonomically affiliated LR sequences were classified as Bacteria, whereas a substantial fraction of SR reads aligned with Archaea (see below). At the family level, the proportion of non-affiliated reads were very similar, ranging from 27% to 52% for SR and 20% to 31% for LR, with a noticeable trend of increasing disparity between LR and SR affiliations with depth (Fig. 3). At the genus level, non-affiliation percentages rose for both LR and SR reads, spanning from 31% to 45% for LR and 27% to 52% for SR, with both sequencing methods demonstrating a gradual lack of affiliation with depth. At the species level, nearly 100% of the reads were classified as non-affiliated, with only a minimal number of reads that could be taxonomically resolved for SR (21 reads), underscoring the limitations of this method, as expected. For LR however, between 39 % (at 5 cm depth) to 59% of all reads remained unaffiliated, which shows a very strong advantage of the LR sequencing in achieving species-level resolution with the SILVA database (Fig. 3).

**Fig. 3.**
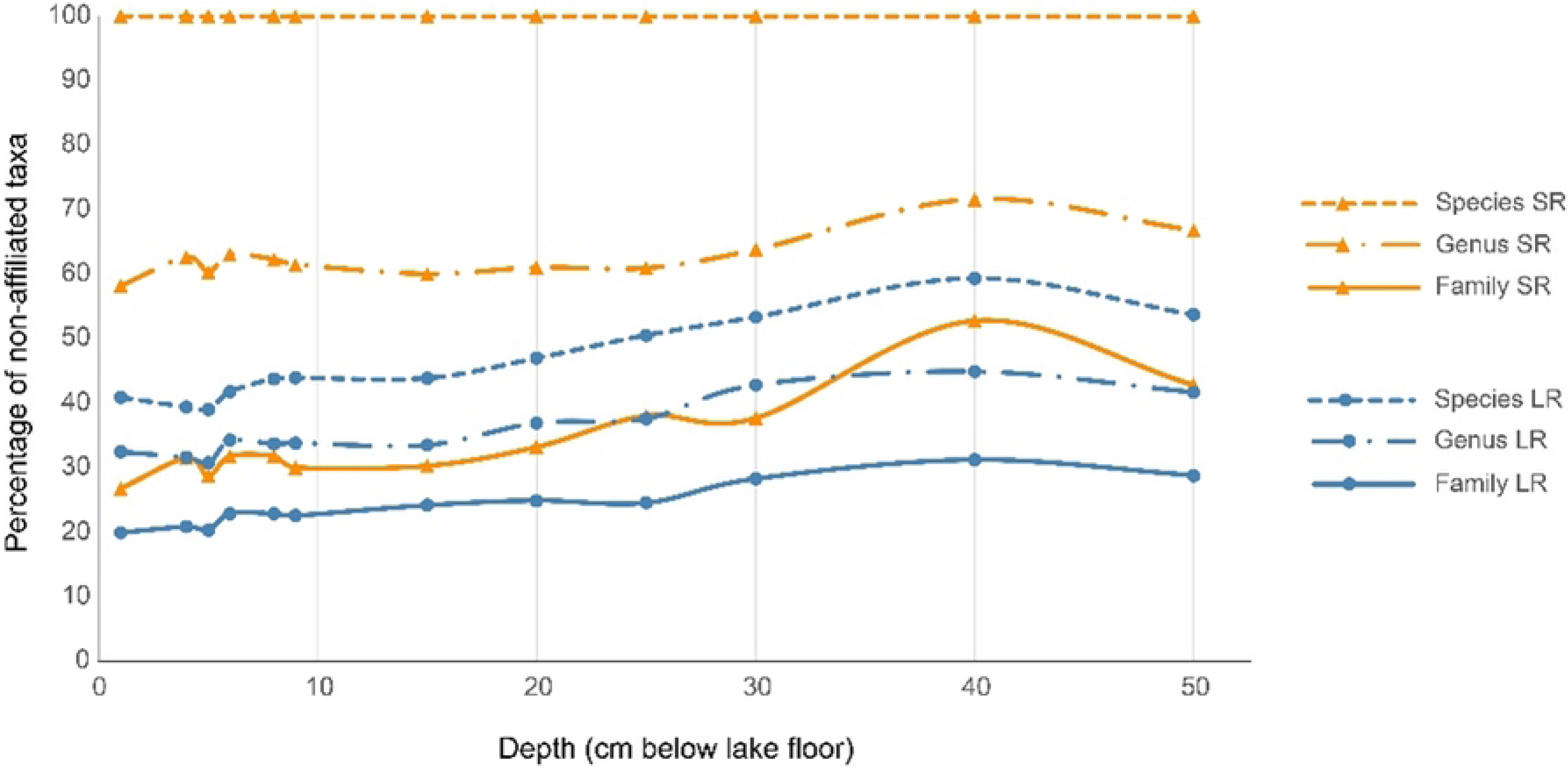
Relative abundance of affiliations at varied taxonomic level. Affiliations are shown at the Family, Genus and Species level, along depth for both SR and LR approaches.

### 4.3. Microbial composition of lake Arnon by sequencing method

For Illumina derived SR, community composition shows high homogeneity across depths, with phyla such as *Actinobacteria, Bacteroidota*, *Proteobacteria*, and to a lesser extent *Desulfobacterota*, exhibiting steady declines in relative abundance with increasing depth (Fig. 4A). Below 20 cm, *Chloroflexi* and *Firmicutes* emerge as dominant phyla. Other phyla, including *Acidobacteria, Patescibacteria*, and *Planctomycetota*, display minimal variation in relative abundance across depths. The PacBio LR sequencing approach shows similar depth-related trends for the major phyla, although with notable variation in relative abundance within samples (Fig. 4B). In the surface samples, *Bacteroidota* and *Proteobacteria* dominate, with *Proteobacteria* comprising an average of 16% of the community (from 29.6 % at 3 cm to 5.0% at 50 cm) and *Bacteroidota* accounting for 17.1% (from 24.4 % at 6 cm to 4.3 % at 40 cm). *Verrucomicrobia* is the third most abundant phylum in surface sample LR-1 (10.3%) but declines with depth, while *Firmicutes* progressively increase in abundance with depth, becoming one of the dominant phyla at deeper layers (8.4% in LR-1, 27.2% in LR-5, and 38.7% in LR-15).

**Fig. 4.**
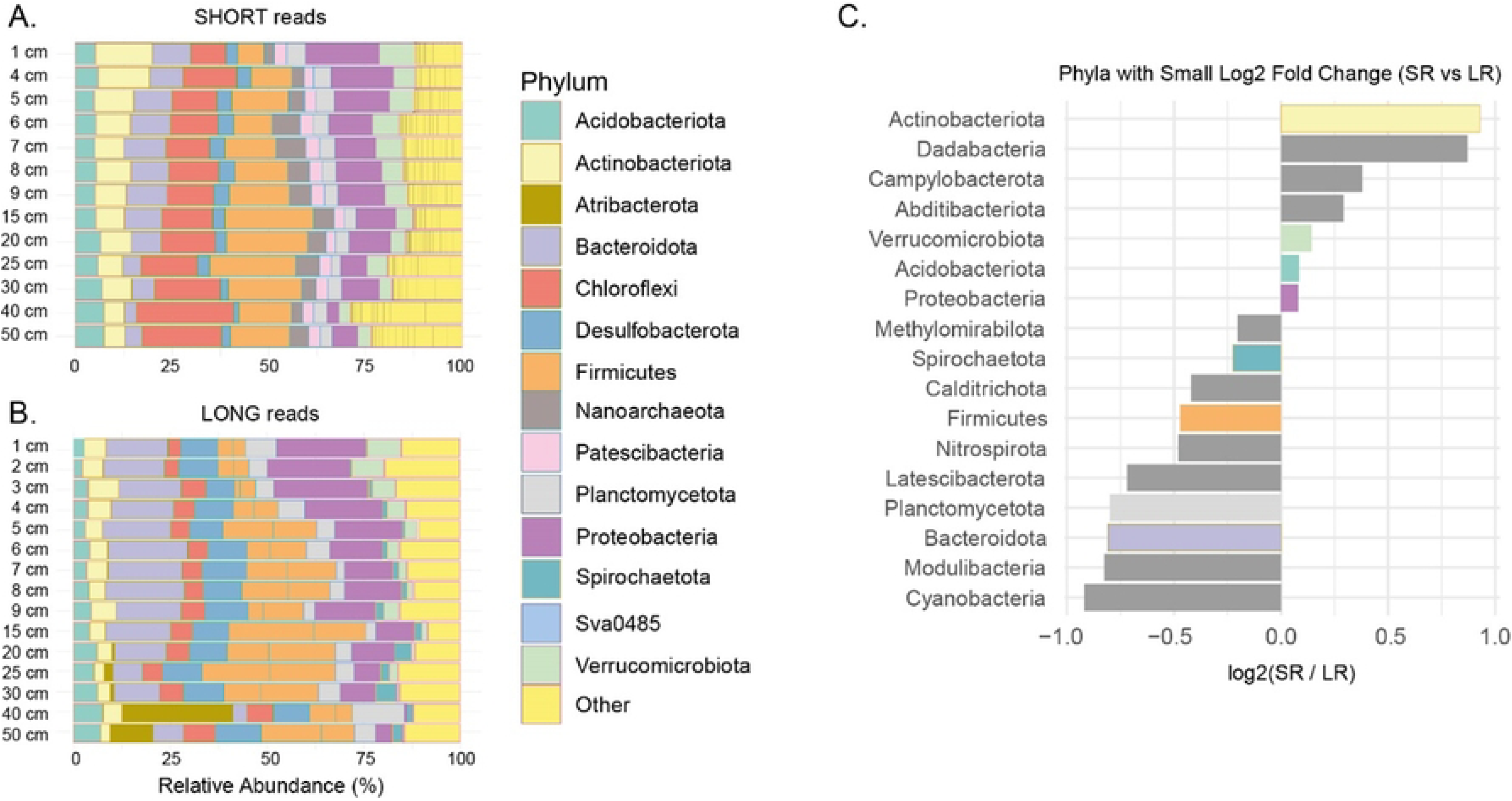
Comparisons of diversity at the phylum level. (A) Barplot of relative abundance of the most abundant phyla for the short read approach. (B) Barplot of relative abundance of the most abundant phyla for the long read approach. For (A) and (B), the remaining taxa are pooled under the “Other” affiliation. (C) Phyla with minimum Log2 Fold changes (< 1) showing phyla with similar relative abundances between LR and SR (phyla appearing in A - B have the same bar color in C).

Long- and short-read sequencing of Lake Arnon samples revealed several key differences in the relative abundance of specific taxa. *Acidobacteria* exhibited notably greater variation with depth in LR sequencing compared to SR. In contrast, *Actinobacteria* (Fig. 4C), *Chloroflexi*, and *Patescibacteria* were consistently lower in abundance in LR sequences, with the latter rarely exceeding 2% and thus categorized under “Other” in LR analyses (Fig. 4). *Nanoarchaeota* were not detected in LR sequencing (only a few reads were assigned to some *Asgardarchaeota* superphylum). Conversely*, Bacteroidota*, *Firmicutes, Desulfobacterota* and *Spirochaetota* showed higher relative abundance in LR than SR data (Fig. 4).

Figure 5 illustrates depth profiles for several taxa of interest in subsurface sediments, highlighting distinct patterns across sequencing methods. *Desulfobacterota* exhibited relatively different depth profiles, increasing from the surface to a peak at 8 cm for SR (4.2%) while relative abundance were three times higher for LR (11.3% at 1 cm to 13.5% at 8 cm). The SR *Desulfobacterota* exhibited a gradual decline to a minimum at 40 cm (SR-40: 1.3%), while for LR the relative abundance remained high and even increased to a maximum at 50 cm (13.8%). *Chloroflexi*-related sequences showed their lowest relative abundance at the surface (1 cm; SR: 9.2%, LR: 3.9%), steadily increasing with depth to reach a maximum at 40 cm for SR (25.0%) and 50 cm for LR (9.7%). *Caldatribacteriota* were nearly absent in the upper sediment layers but became significant at 40 cm, reaching 9.5% in SR and 32.6% in LR. Taxa from *Spirochaeta*, detected exclusively in LR data, also increased with depth, from 0.5% at 1 cm to a peak of 6.6% at 30 cm. Archaeal sequences, detectable only through SR except for a few *Asgardarchaeota* in the top layers for LR, displayed a peak at SR-6 cm, followed by a decline with depth and additional increases at 25, 40, and 50 cm, represented by a diversity of archaeal taxa Fig. 6).

**Fig. 5.**
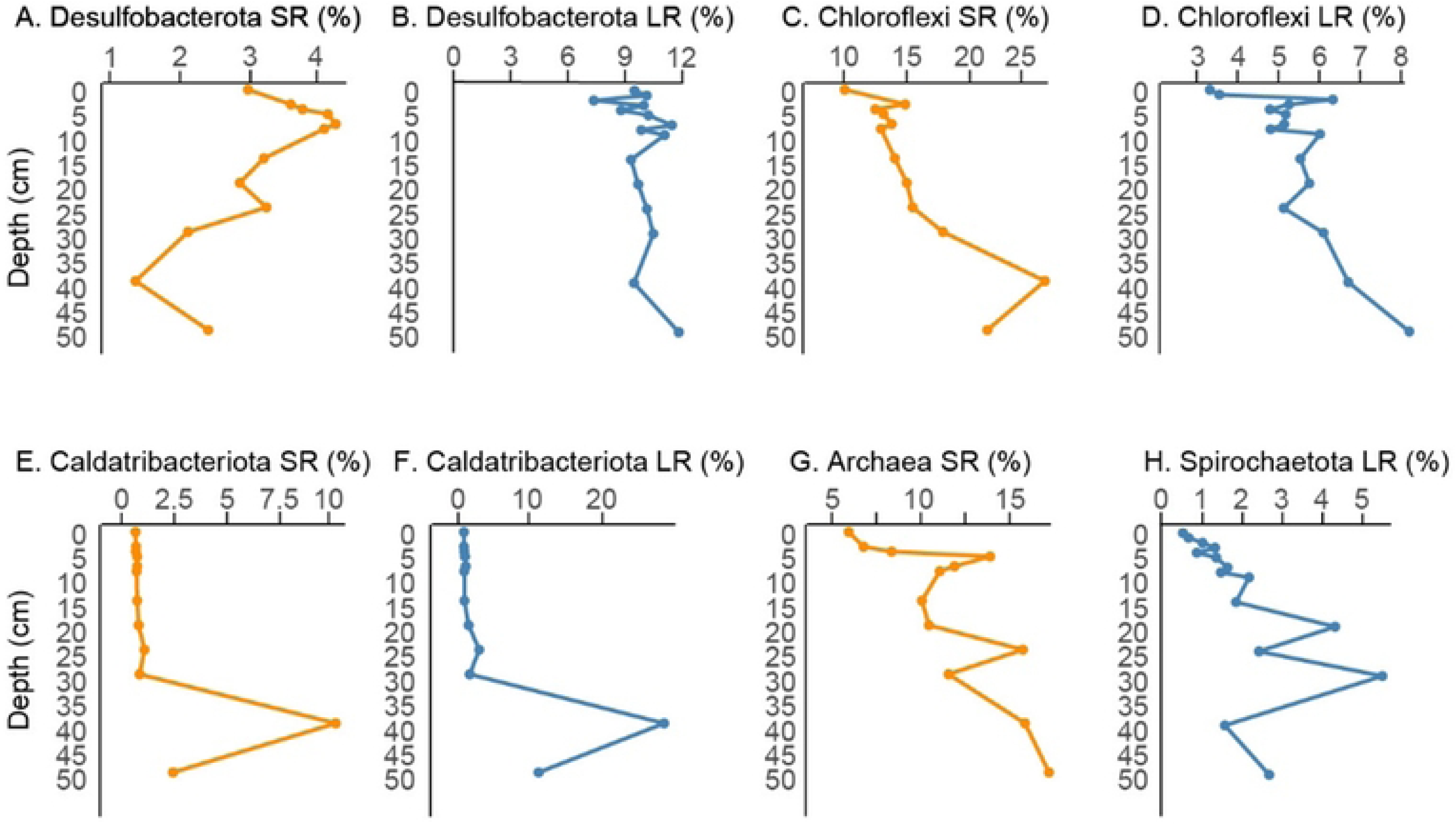
Relative-abundance of key taxa along depth for different sequencing approaches. *Archaea* were not detected below 2 cm for LR. *Spirochaeta* represented a minor portion of the taxa in SR. Orange codes for SR, blue for LR.

**Fig. 6.**
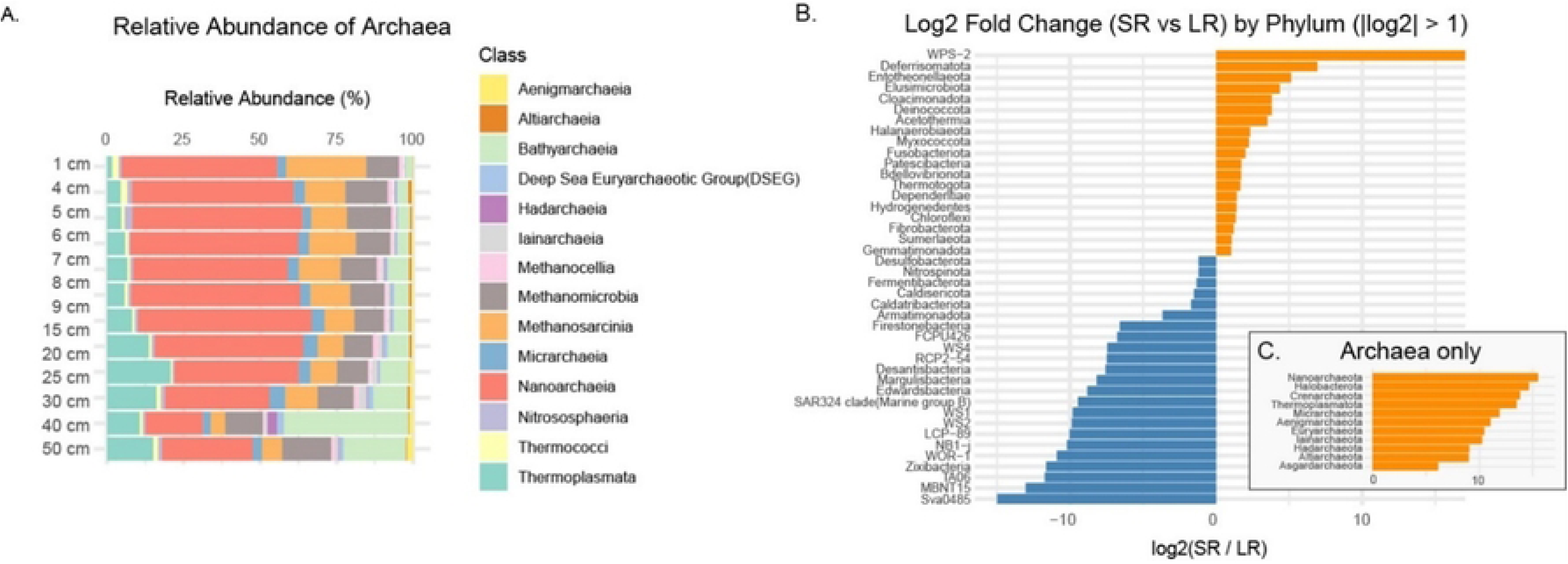
Archaeal abundance for SR and comparison of rare taxa and archaea with LR. (A) Barplot of the relative abundance of Archaea classes in the Short Read pool, and maximum log2 fold change between SR and LR (>1) for bacterial (B) and archaeal phyla (C).

### 4.4. Archaeal Recovery and major differences in taxonomic affiliations

Despite employing relaxed parameters in the pb16Snf pipeline, very few archaeal sequences were identified in the PacBio dataset when compared to the SILVA database. Lowering the VSEARCH identity threshold to 0.7 and adjusting error filtering parameters increased the total number of ASVs classified but did not improve the recovery of archaeal taxa. The composition of *Archaea* is therefore only investigated using SR. At the class level, *Nanoarchaeia* dominated the levels above 30 cm (Fig. 6A), although a steady decrease in relative abundance was observed with depth. *Thermoplasmata* and to a lower extent *Bathyarchaeia* increased gradually with depth. *Bathyarchaeia* became the dominant class at 40 cm, and still held ca. 20% of the relative abundance at 50 cm, similarly to *Nanoarchaeia*. Methanogenic taxa, including *Methanocella*, *Methanosarcinia* and *Methanomicrobia*, formed a substantial share of the total abundance (30% for LR1), and decreased gradually to 20% altogether with depth (with *Methanomicrobia* slowly taking over *Methanosarcinia* as the main methanogenic taxon). *Methanomicrobia* is the 4^th^ most abundant taxa after *Thermoplasmata* in LR-50. But the overall relative abundance of *Archaea* increases with depth (Fig. 5G).

Overall, archaeal phyla and *WPS-2* and *Deferrimonadota* phyla had log_2_-fold changes superior to 5, in favor of SR, while *Sva0485, MBNT15, TA06*, *Zixibacteria*, *WOR-1* and *NB1-j* were over-represented in LR (Fig. 6B-C).

### 4.5. Betadiversity and community structure

PERMANOVA analysis revealed that both the sequencing method (SR vs LR) and sediment depth had significant effects on microbial community composition (Bray-Curtis dissimilarity; *p* = 0.001 for both factors; NMDS visualization available in the supplementary material Fig. S1). The sequencing method explained approximately 29.7% of the total variance (R² = 0.297), while depth accounted for 35.6% (R² = 0.356), indicating that depth was the stronger predictor of community structure in this dataset.

Species contribution to betadiversity was measured for both techniques. A number of ASV with similar affiliations were found in both data set to carry an important share of the SCBD. Notably, *Caldatribacterota JS1, Bacteroidia Vadin17, Desulfobacterota, Verrucomicrobiota Luteolibacter* and *Cyanobacteria* (Fig. 7). However, numerous differences were also identified as SCBD top contributors, for example *Xanthomonadaceae*, *Chloroflexi*, *Spirochaeta*, and methanogenic archaea along with other archaeal clades *Bathyarchaeia*, *Methanomassiliicoccales*, *Halobacterota Methanosaeta* or *Woesarchaeales* for SR, and *Firmicutes* ASV, *Bacteroidota Ignavibacterales*, *Sva0485* or *MBNT15* (Fig. 6B, 7B) for LR.

**Fig. 7.**
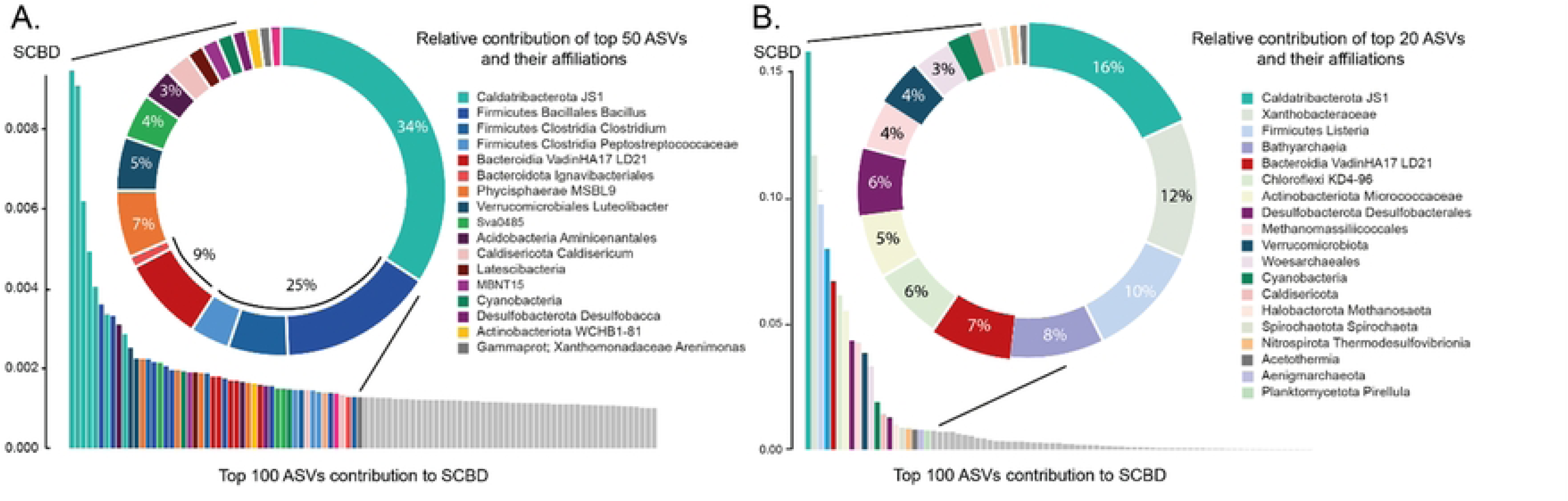
Species contribution to beta-diversity. (A) SCBD from the top 100 ASV and affiliation of the first contributors for LR. (B) SCBD from the top 100 ASV and affiliation of the first contributors for SR. Corresponding affiliations in SR and LR are coded with similar colors.

Distinct structural differences emerged between the LR and SR co-occurrence networks (Fig. 8), despite both displaying a broadly similar architecture organized into four major modules (modularity of 0.54 for SR and 0.49 for LR, Table S3). Mean degree (the average number of connections per ASV) was 30.21 for SR and 42.81 for LR, with a higher clustering coefficient for LR (0.71) than SR (0.52). Betweenness centralizations were quite similar, (SR : 0.02and LR : 0.016), with higher degree for LR (0.116) compared to 0.079 for SR. In both datasets, the dominant module (Module 1 in SR, Module 2 in LR) consisted of a large number of ASVs exhibiting relatively moderate degrees of connectivity (Fig. 8C-D). A second prominent module accounted for 26.5% of the network in SR (module 2) and 41% in LR. In SR it was composed of ASVs with higher individual degrees, while mean degree was quite similar for module 1 and 2 for LR, where it was module 4 (13,4%) that had higher mean degree, and lower Simpson dominance (0.75, Table S3).

**Fig. 8.**
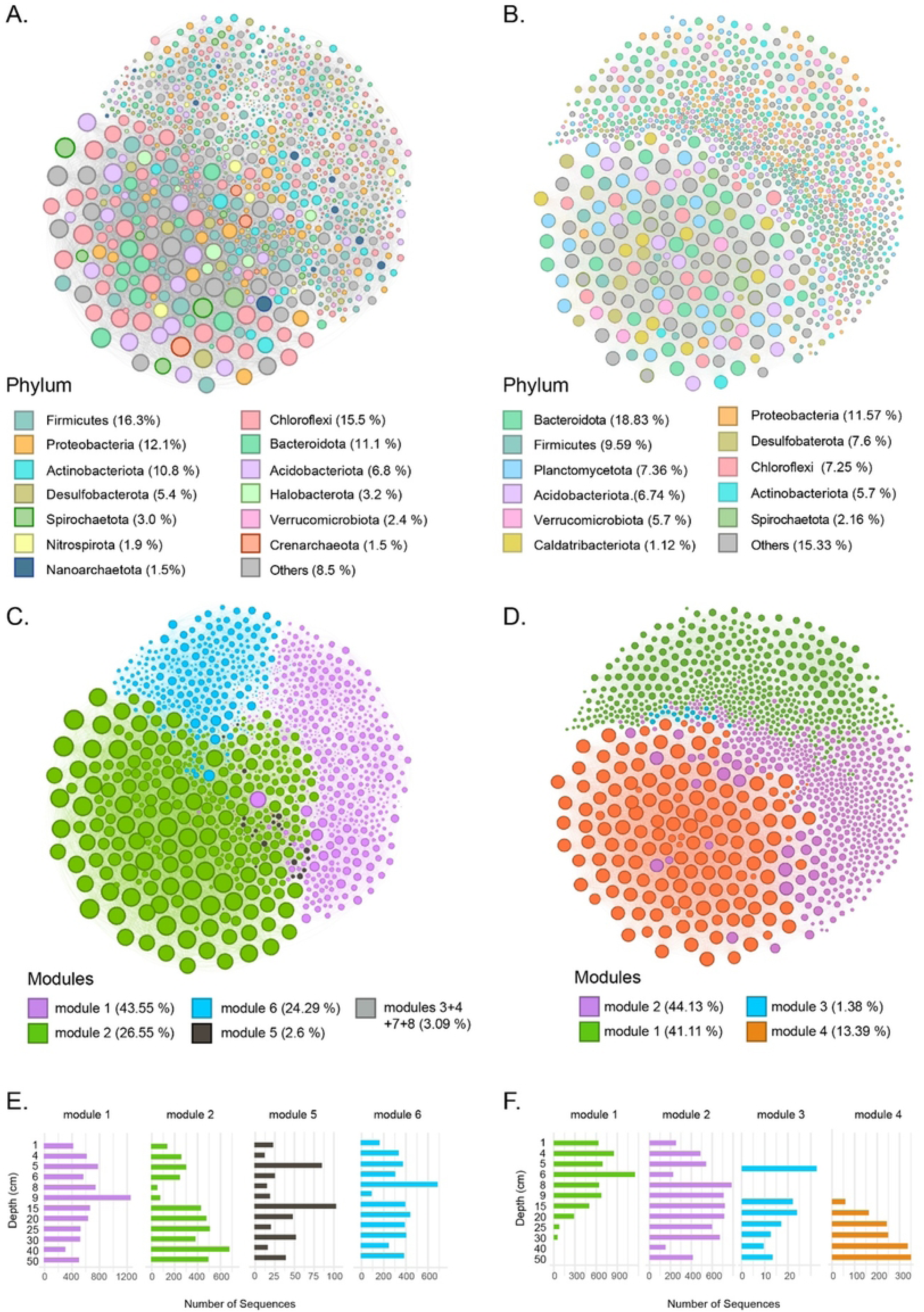
Co-occurrence networks of the microbial community sediment samples. Co-occurrence networks with ASVs colored by taxonomy for SR (A) and LR (B), and by modularity for SR (C) and LR (D). The size of each node is proportional to the number of connections. Total number of sequences at each sampling depth for each ASV in the main modules for SR (E) and LR (F).

In the LR dataset, module 4 was primarily associated with ASVs from the deeper sediment layers. Module 2 in the SR dataset was also more represented in deeper layers (below 15 cm), but with abundant reads also above, showing a difference in modularity structure.,. A fourth module (2.6 % for SR and 1.38% for LR) while occupying a similar position within the network topology in both datasets, displayed differing depth distributions between LR and SR (Fig. 8C-D). When nodes were colored by phylum-level taxonomy, the network modularity did not show marked patterns.

## 5. Discussion

### 5.1. Microbial diversity of Lake Arnon

The analysis of 16S rRNA gene sequences from Lake Arnon sediments revealed a diverse microbial community, exhibiting transitional shifts across sediment depth without a significant loss of diversity (Fig. C). At the phylum level, *Proteobacteria*, *Bacteroidota*, *Firmicutes*, *Acidobacteria*, *Actinobacteria*, *Chloroflexi*, *Planctomycetota*, and *Patescibacteria* were predominant throughout the sediment core, consistent with observations from other lakes of similar depth. (e.g. Berg et al., 2022; Han et al., 2020; Pearman et al., 2021).

Several taxa displayed depth-related changes, reflecting patterns commonly observed in sedimentary environments. This was particularly evident for *Chloroflexi*, which increases in relative abundance with depth, reaching its maximum at 40 cm in the SR dataset and 50 cm in the LR dataset. *Chloroflexi* were consistently present throughout the sediment core, with specific lineages showing distinct vertical trends. Genera such as *Anaerolineae*, the *GIF9* group, and the *csp1-4* clade were detected across the entire core, with members of the *GIF9* lineage becoming dominant at greater depths (Supplementary Table). These groups have been reported in various anoxic environments, ranging from continental to oceanic settings, and are associated with metabolic strategies suited to deep, low-energy conditions—including fermentation of a broad spectrum of substrates and (homo)acetogenesis (40–42). *Spirochaetota*, whose relative abundance increased with depth, is commonly found across a range of environments, including freshwater, methane-rich, and hypersaline sediments (43–46). Showing a similar depth-related distribution, *Caldatribacterota,* a lineage largely specific to deep sedimentary systems (6,47), exhibited a sharp increase in abundance at and below 40 cm in both the LR and SR datasets (Fig.5E-F). This pronounced shift appears to be specific to our sedimentary record and plays a key role in structuring microbial beta diversity, as indicated by the high Species Contribution to Beta Diversity (SCBD) values associated with *Caldatribacterota JS1* ASVs (Fig. 7).

Metagenomic and metatranscriptomic studies have shown that *Caldatribacterota* possess highly specialized gene repertoires adapted to low-energy environments, including pathways for fermentation and potentially acetogenic metabolisms (47,48) for members of this phylum. Their emergence below 40 cm may represent a geochemical or energetic boundary, beyond which taxa specialized for life under organic resource limitation begin to dominate. This is further supported by the concurrent increase in *Chloroflexi* and various archaeal lineages in the SR dataset (Fig. 5-6).

Figure 7 reveals that several archaeal ASVs contributed significantly to the SCBD. Notably, the increasing abundance of *Bathyarchaeia*—which peaked around 40 cm (Fig. 6)—coincided with a decline in sequences affiliated with *Methanosarcinia* and *Nanoarchaeia*. This shift may reflect a transition to the utilization of more complex organic carbon sources (49–51), or a marked shift to acetogenesis (52,53).

The distribution pattern of *Desulfobacterota* in the sediment column of Lake Arnon is not straightforward. Although sulfur cycling was evident through precipitation of diagenetic sulfate and sulfur minerals such as pyrite and barite in the sediment (identified but to be detailed in a forthcoming study; Laakkonen et al., in prep), the relative abundance of *Desulfobacterota* ASVs differed notably between sequencing approaches. In the LR dataset, their abundance was relatively high and stable below 10 cm, fluctuating between 9% and 10%, with a maximum of approximately 12% at 50 cm. In contrast, the SR dataset showed a declining trend with depth, from about 4% at 10 cm to below 2% at 40 cm. At greater depths, the dominant *Desulfobacterota* ASVs were affiliated with the uncultured *BSN033* lineage (54). Although metagenome-assembled genomes (MAGs) from this group have been reported to lack the canonical *dsrAB* genes typically associated with dissimilatory sulfate reduction, they possess a suite of genes involved in anaerobic hydrocarbon degradation, including those encoding enzymes for the breakdown of aromatic compounds. At shallower depths (above 15 cm), *Desulfobaccia*, *Desulfobacteria*, *Syntrophia*, *Syntrophorhabdia*, and *MBNT15* were more abundant, while *BSN033* remained present. Genomic reconstructions from organic-rich, cold peatlands have shown that the *MBNT15* clade comprises versatile facultative anaerobes capable of dissimilatory iron reduction (55). *Desulfobaccia* and *Desulfobacteria* possess *dsr* genes associated with dissimilatory sulfate reduction (56).

Interestingly, the co-occurrence of MAGs assigned to *Desulfobaccia*, *Syntrophia*, and *BSN033* has also been reported in the low-sulfur sediments of Lake Superior (57), where sulfur cycling appears to depend on a diverse microbial community capable of reducing organic sulfur compounds. In our dataset, ASVs associated with *Desulfobacterota* members of the class *Syntrophia* were closely related to species such as *Smithella* and *Syntrophus aciditrophicus*. Representatives of these taxa have been shown to degrade long-chain alkanes, fatty acids, and benzoate through syntrophic interactions with hydrogenotrophic methanogens (58,59).

Collectively, the microbial community composition of Lake Arnon sediments reflects a complex network of metabolic interactions, with depth-related shifts corresponding to changes in substrate availability. These findings are consistent with studies from other alpine and freshwater lake sediments, highlighting the influence of environmental gradients on microbial distribution and function. Inference on metabolic potential can be pushed a little further thanks to species-level association powered by the PacBio LR approach. The SR approach reveals archaeal groups with key diversity and metabolic roles in the sediment subsurface, which are overlooked by LR. The LR approach allowed the detection and identification of rare taxa that have significant importance in sediment environments. It is the case of members of the *sva0485* taxon, identified in numerous sediment settings, in association with anaerobic methanotrophs (60) and/or metal contaminated sediments where there were suggested to cycle iron and/or sulfur (61,62). Other uncultured clades like *Zixibateria, TA06* or *WOR-1* were preferentially detected by LR and are alleged actors of C, N and S cycling in sedimentary anoxic environments (63,64).

### 5.2. Lack of Archaeal Sequence Detection in Long Reads

Pacific Biosciences (PacBio) long-read sequencing has significantly improved its initial error rates, making it an increasingly reliable and affordable approach in microbial ecology (65,66). It now generates long reads (>10 kb) with very high accuracy following circular consensus sequencing (CCS) error correction. This approach offers a major advantage in resolving full-length 16S rRNA genes. The improved resolution allows for accurate taxonomic classification at the species and even strain levels, reducing biases associated with sequence reconstruction from short fragments—whether in MAG-based approaches or metabarcoding diversity studies. Long-read sequencing is particularly beneficial for identifying novel or closely related taxa that are often misclassified or remain unresolved with short-read methods. However, because such high accuracy is relatively recent, only a limited number of studies have applied this approach to sedimentary environments. The results of our study demonstrate that long-read sequencing provides a clear improvement in species-level resolution, enabling more detailed insights into microbial metabolic potential and community structure (Fig. 3 and Fig. 7). Using the same reference database (SILVA nr99, version 138.1), phylogenetic assignment based on LR sequencing proved significantly more effective, allowing species-level affiliation for over 50% of the obtained ASVs—compared to almost none with short-read (SR) sequencing. At the genus level, more than 60% of ASVs could be assigned to known taxa with LR, while SR yielded affiliations for fewer than 40% of ASVs. However, in the context of subsurface studies, taxonomic affiliation alone does not necessarily translate to an understanding of functional potential. Functional inference is generally achieved and validated through culture-based experiments—an approach that finds strong limitations for subsurface studies (67). The deep biosphere is estimated to contain up to 80% of uncultured microbial species (Lloyd et al., 2018), whose functions can only be inferred through genomic reconstructions. Many of the sequences present in current databases originate from metagenome-assembled genomes (MAGs), which are often incomplete and potentially contaminated (Setubal, 2021). As such, much of the so-called "microbial dark matter"—which is of particular interest in this study—relies on functional interpretations derived from these MAGs (Lloyd et al., 2018).

The PacBio LR metabarcoding approach clearly provides valuable insights, as demonstrated above, but it also presents certain limitations. One notable limitation is the insufficient coverage of archaeal species. This is evident in our dataset, where Archaea constitute a consistent 10– 15% of the community based on SR data, yet are barely detectable in the LR dataset, even when using relaxed parameters during post-sequencing analysis.

Archaeal dominance is well-documented in deep sedimentary environments, both marine and lacustrine, highlighting the importance of adequately capturing this domain in microbial community studies (7,68). For example, BLAST analysis of the LR dataset at 25 cm against the SILVA database revealed sequences with best hits to Archaea, though with low identity scores (∼79–81%) and a high number of mismatches. Additionally, these putative archaeal sequences did not represent more than 15% of the total reads, in contrast to the proportions observed in the SR dataset. When archaeal ASVs identified in the SR dataset were blasted against the PacBio reads from the same sample, approximately 25% of the reads produced matches, but with generally poor sequence identity (∼75–80%), suggesting either low archaeal abundance or substantial sequence divergence from known archaeal references. Further BLAST searches against the NCBI database yielded similarly low identity values (∼70–80%) for these putative archaeal sequences, supporting divergent archaeal lineages, that could be specific to this sedimentary environment, or overall database limitations (common for Archaea).

One possible explanation is that the primers used in the PacBio full-length 16S rRNA protocol did not efficiently amplify archaeal sequences. This hypothesis is supported by results from our community controls (Fig. S2). Although *Euryarchaeota* members such as *Methanosaeta* and *Methanoregula* were not listed in the manufacturer’s Zymo Community Control dataset, they were detected in the SR dataset—yet remained undetected in the LR dataset. Previous studies have shown that so-called universal primers often have differential amplification efficiencies across bacterial and archaeal taxa, which can lead to the underrepresentation or complete omission of certain groups (17). This limitation was also recently highlighted for primers used in long-read sequencing of the 16S–ITS–23S operon (21).

The absence of archaeal sequences in the LR dataset may also be partly attributed to the poor quality of archaeal DNA in the control samples, given that these taxa were not expected to be present initially. Long-read sequencing technologies, such as PacBio, require high-quality, high-molecular-weight DNA for optimal performance (66), and it is possible that archaeal DNA fragments were too degraded or too short to be effectively amplified. This challenge is especially relevant for sedimentary samples, where DNA degradation is common due to prolonged burial and exposure to environmental stressors (Willerslev et al., 2004). Archaeal DNA in the sediment samples may also be degraded or fragmented to a degree that hinders amplification using the PacBio protocol. This possibility is modestly supported by the decreasing LR-to-SR ratio with depth (Fig. 2D), although additional samples would be needed to establish statistical significance. However, to our knowledge, there is no evidence in the literature suggesting preferential degradation of archaeal versus bacterial biomass in sedimentary environments. In fact, existing studies suggest that all microbial biomass is subject to recycling, despite the greater resistance of archaeal membranes to extreme environmental conditions. (69–71).

Even if archaeal sequences are present in the samples, they may be at such low abundance that they do not pass the denoising and filtering steps of the pipeline. However, considering the significant presence of *Archaea* in comparable sedimentary environments (37,68,72,73) and their detection at relatively high levels in the short-read dataset (reaching up to 15% in the deeper layers), it is unlikely that their absence in the long-read dataset reflects a true biological pattern. Interestingly, PacBio sequencing appears to be less affected by variation in relative abundance, as shown by results from mock community samples. Besides, the relative abundance of major taxa detected in the PacBio LR Zymo community control closely matched the manufacturer’s documented composition (ZymoBIOMICS Microbial Community Standard 16S rRNA Gene Abundance: https://files.zymoresearch.com/pdf/d6300-_zymobiomics_microbial_community_standard_v1-1-3.pdf), whereas the SR data show more divergent relative abundances, with a notable bias toward *Lactobacillus*.

The absence of archaeal sequences in the PacBio full-length 16S rRNA dataset raises several hypotheses that warrant further investigation. While it is widely accepted that sedimentary environments host diverse and abundant archaeal communities (e.g., Lloyd et al., 2013; Orsi et al., 2020), our analysis suggests that their apparent absence in the long-read dataset is more likely due to methodological limitations than to a true biological absence. Specifically, the limited detection of Archaea in the LR data appears to be primarily related to primer selection and database biases against longer archaeal sequences, rather than an overestimation of archaeal abundance in the SR dataset.

### 5.3. Meaningful interactions vs rare diversity

Another point that needs to be addressed is the differences obtained from betadiversity analysis we performed between SR and LR. While sediment depth remained the primary driver of community differentiation, sequencing method introduced a substantial bias in microbial profiles. On the SR side, the species contribution to betadiversity was carried by a phylogenetically diverse number of ASV, ranging from *Caldatribacterota*, to *Bathyarchaeia, Firmicutes, Desulfobacterota* and other archaeal ASV. Network analysis of SR also showed higher modularity and module numbers, including small modules with low richness. When analyzed with the LR approach, *Caldatribacterota* and *Firmicutes* carried most of the SCBD, and even though modularity was relatively similar, more structuration (depth-based) was observed within modules.

The observed discrepancies in beta diversity structure and network modularity between the SR and LR approaches can be explained by a set of interrelated hypotheses, stemming from the sequencing depths, taxonomic precision, and ecological signal retention between the two methods:

LR offered a more centralized, cohesive and taxonomically organized view, likely due to better sequence resolution and affiliations. Larger modularity and fragmentation for SR, even from a rarefied dataset shows more noise and capture of rarer taxa. Network differences remain therefore intrinsic to the sequencing method.

LR-based networks exhibit modules that are phylogenetically diverse and compositionally even, implying that network structure is driven less by taxonomy and more by shared ecological roles or environmental interactions. A vision that is likely to reflect the reality of complex ecosystem. Lower fragmentation, dominance and evenness in LR modules compared to SR modules is possibly linked to the improved resolution of full-length 16S sequences, which allows ecologically similar but taxonomically diverse ASVs to be correctly grouped within the same module. In contrast, SR modules may artificially cluster ASVs from the same broad taxonomic lineage, inflating the dominance of particular phyla (Fig. 8A-B) (74,75). The absence of archaeal sequences in LR may also have strong impact on our network visualization, missing an important feature of the sediment diversity, and simplifying it to a point that allows to visualize depth-related modularity for example.

## 6. Conclusions

In the study of deep biosphere microbial communities, sequencing technologies play a pivotal role in accurately characterizing the complex assemblages inhabiting sedimentary environments, even for 16S profiling. PacBio LR sequencing offers notable advantages, particularly through its ability to resolve full-length 16S rRNA genes, thereby enhancing taxonomic classification down to the species or even strain level. This high-resolution capability is especially valuable in deeper sediment layers, where clades such as *Caldatribacterota*— associated with specialized fermentative metabolisms under anoxic conditions—become more prevalent and can be reliably linked to known MAGs. In Lake Arnon, this approach allowed us to identify a shift toward the utilization of more specific organic substrates, including indications of homoacetogenic activity. Additionally, beta diversity analyses based on LR data revealed patterns that likely reflect more ecologically meaningful interactions within the subsurface microbial community.

Despite these strengths, the LR approach also revealed important limitation, chief among them being the choice of primers. While it can be easily resolved with more applications and experienced choices, the underrepresentation of key archaeal clades in this study severely constrained our understanding of the subsurface biosphere ecology. Taxa such as *Halobacterota* and *Bathyarchaeia*, known to play central roles in methanogenesis and the degradation of complex organic matter, are essential contributors to carbon, nitrogen, and sulfur cycling in sedimentary environments. These groups were identified in the SR dataset as key drivers of community structure and diversity, yet were largely absent from the LR data, leaving our ecological interpretation incomplete.

When addressing questions related to the deep biosphere, it is important to acknowledge that metabarcoding approaches targeting specific marker genes such as 16S rRNA provide limited insights into microbial functional potential. In contrast, shotgun metagenomic sequencing offers a more comprehensive view, capturing the entire genomic repertoire of the community, including functional genes and metabolic pathways. With ongoing improvements in long-read technologies—such as PacBio’s HiFi reads, which combine length with high accuracy—we are now approaching a more realistic representation of ecological and functional interactions in the subsurface. However, the inclusion of archaeal diversity remains critical and requires primer sets specifically designed to capture this domain. With such methodological refinements, the depth and affordability of SR sequencing may soon be matched by LR technologies, marking a major step forward in our ability to study the microbial ecology of these poorly understood environments.

## Supplementary materials

Fig. S1 : NMDS of Lake Arnon samples (Bray-Curtis distance) color-coded by sequencing method at the phylum level, after rarefaction to total LR reads.

Fig. S2 : taxonomic affiliation of both controls for Sr and LR. Note the detection of archaeal sequences (black color) in the SR controls.

Table S1 : Total abundance per sample and complete taxonomic affiliations of ASVs detected using the long read approach

Table S2 : Species contribution to betadiversity for LR and SR, with their corresponding affiliations.

Table S3 : network metrics and module wise diversity indexes for SR and LR.

## Acknowledgement

We thank Gaetan Sauter for his assistance during fieldwork, and Simone Oberhänsli for her biostatistical work and constructive suggestions, as well as Pamela Nicholson for suggestions and fruitful discussions on the strategy and results.

During the preparation of this work the author used ChatGPT to curate and correct their R scripts, and to enhance the English and the text structure and readability. After using this tool/service, the author reviewed and edited the content as needed and takes full responsibility for the content of the publication.

